# Reducing the environmental footprint of laboratory research using visual communications: A pilot study in Spain

**DOI:** 10.1101/2025.09.11.675379

**Authors:** Raquel Marco-Ferreres, Silvia Ayora, Carmela Cela, Álvaro Sahún-Español, María López-Sanz, Pablo Izquierdo

## Abstract

Addressing the environmental impact of laboratory research provides a meaningful opportunity for workplace-level mitigation. However, in Spain, the importance of sustainability in research is only starting to be recognised at a national level. We developed a scalable, visual-oriented communications strategy that was designed with input from scientific staff and included a live seminar and a coherent set of graphic materials. Five pillars were targeted: three key areas of consumption (energy, water, and plastic use), procurement, and colleague engagement. The scheme was piloted in two of the largest biomedical research centres in Spain: the Severo Ochoa Molecular Biology Centre (CBM) and the Spanish National Centre for Biotechnology (CNB). The pilot engaged 50 laboratories and services across both centres, of which 34 were followed up for 5 months. The results indicate engagement with sustainability measures and shifts toward best practice across metrics, including equipment switch-off, water use, plastic item reuse, and purchasing decisions. During the follow-up period, 23% of participants adopted new measures related to freezer maintenance, decluttering, or sample organization, 55% removed old data from storage to save energy, 100% reduced their use of lifts, and 94% discussed research sustainability with their laboratory/group. The findings also revealed key areas for improvement, including the management of shared resources and resistance to change protocols. These results offer a wide range of quantitative and qualitative insights to guide further efforts and scaling at regional and national levels.

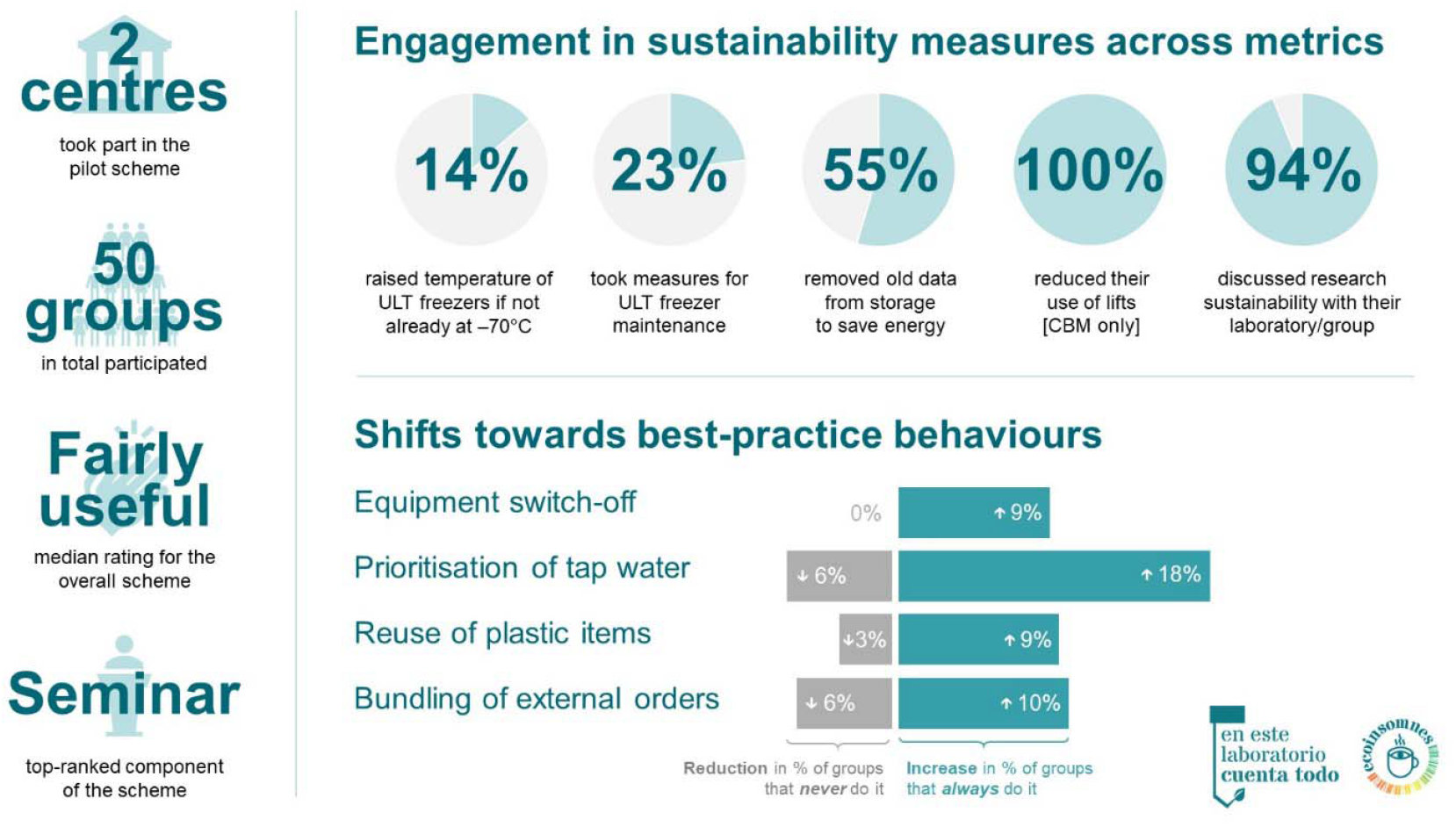

**Sustainability spotlight:** This paper describes a scalable, visual-oriented communications strategy that can be used to promote sustainable behaviours in research laboratories. We present pilot results indicating engagement in sustainability and shifts toward best practice across metrics. Our work contributes to UN Sustainable Development Goal (SDG) 9 (industry and infrastructure) and SDG 12 (responsible consumption and production).

## Introduction

The current climate emergency, caused by the burning of fossil fuels, is rapidly unfolding globally (1). As a result, Europe has experienced record-breaking heat, droughts, and floods in recent years (1,2). The climate crisis is also increasing the frequency and intensity of wildfires, climate-driven spread of pathogens and disease vectors, and heat-related deaths across the continent (2). Without mitigation, adverse climate-related health impacts are expected to worsen worldwide, affecting billions of people (2).

Research has a non-neglectable climate and environmental impact, which presents a valuable opportunity for mitigation within the workplace. Research laboratories use from 3 to 10 times more energy than an equivalent-size office (3,4). That energy demand has a significant associated carbon footprint; although the associated CO_2_ emissions depend on each country’s electricity mix, emissions can reach 5 tonnes per year for an ultra-low temperature freezer (ULT) set at –80°C (5) and 14 tonnes for a fume hood (6). Research laboratories are also responsible for an estimated 5 million tonnes of plastic waste each year (7,8), and high volumes of pipette tips, pipettes, and gloves are used daily (9).

However, there are multiple simple measures that have been shown to be effective. For example, closing the sash of a fume hood can save ~40% of energy in addition to being safer (10,11). Raising the set temperature of ULT freezers from –80°C to –70°C can save ~28–36% of energy (12,13) (critically, without affecting sample safety or integrity for many applications (14). Removing old samples to avoid clutter, removing ice build-up and defrosting regularly, and keeping up-to-date freezer maps or inventories to save time finding samples may also contribute to energy saving. With regards to the energy demand from computational research and the storage of digital data (15,16), periodic removal of old/obsolete data (including large image, videos, and sequencing files) can be an effective strategy, particularly as the energy use associated with storage is expected to rise as more data are generated. Resource efficiency can also be improved for water use. For instance, using distilled water instead of milli-Q water, or tap water instead of distilled water when feasible, can substantially reduce water demand. Producing one litre of distilled water requires approximately three litres of input water, while one litre of milli-Q water requires roughly five litres (12). Finally, the use of single-use laboratory plastics can be significantly reduced by replacing disposable plastic items with autoclavable glass alternatives, adopting best practices to minimise pipette tip use, and reusing gloves or weighing boats when appropriate (17).

Notably, a large-scale international survey conducted by the Royal Society of Chemistry (RSC) showed that 79% of researchers are aware of the environmental impact of their work, but only 63% have taken steps to mitigate it, highlighting a 16-point gap between awareness and action across the research community (10). Multiple initiatives have been rolled out in different countries to promote sustainable research practices; the Laboratory Efficiency Assessment Framework (LEAF) (18) and the My Green Lab (19) scheme are landmark examples. A vast array of guidance, best-practice documents, and visual aid are currently available from several organizations, mostly in English-speaking countries and Germany (17,20).

In Spain, current efforts toward sustainable research are led at a local or regional level (e.g., Catalonia’s SuRe-Cat workgroup) (21). However, the importance of sustainability in research is only starting to be recognised at a national level, with the Spanish National Research Council (CSIC; the country’s leading research institution) presenting its first ever sustainability plan in May 2025 (22). Critically, the RSC survey highlighted lack of training and time constraints among the top challenges that researchers find to implement sustainability measures in their laboratories (10). In this context, we conducted a pilot study in two large CSIC research centres in Madrid with the aim to develop a scalable, visual-oriented communication strategy that raises awareness and promotes the adoption of simple yet effective sustainability practices.

## Methods

### Participants

This pilot study targeted research laboratories, scientific services (e.g., microscopy and cell culture), and non-scientific departments (e.g., communications) in two research centres in Spain: the Severo Ochoa Molecular Biology Centre (CBM) and the Spanish National Centre for Biotechnology (CNB). CBM is a joint centre of the CSIC and Universidad Autónoma de Madrid and hosts 97 laboratories and service units. CNB is a CSIC-affiliated centre and hosts 92 laboratories and service units. Both centres have established sustainability committees (formed in 2023 at CBM and in 2022 at CNB), each comprising 17 members representing staff across job roles and departments. At CBM, each laboratory/group has a designated sustainability contact for direct communication with the sustainability committee.

### Development of materials

Five main pillars were targeted: three key areas of consumption (energy, water, and plastic use), procurement, and colleague engagement. With these in mind, a four-way visual-oriented communications strategy (“En este laboratorio cuenta todo” [“In this laboratory, everything counts”]) was developed through a co-creation process that incorporated input from the sustainability committees of both participating centres.

First, we designed stickers to promote the adoption of new measures or the maintenance of pre-existing ones. We focused on actions that were deemed highly effective. Most stickers were designed to be widely applicable, although some varied between settings (e.g., not all laboratories had ULTs or fume hoods) (**Fig. 1A–F**). A subset of stickers targeted communal equipment/areas (i.e., autoclaves, printers, and recycling bins) (**Fig. 1G–I**). Actions for leadership (e.g., provision of renewable energy, heating regulations, or equipment retrofitting) and wider workplace measures (e.g., regarding commuting options, conference travelling, or plant-based choices in the canteen), while of great potential (23), were considered out of scope for this bottom-up, pilot scheme.

**Fig. 1.**
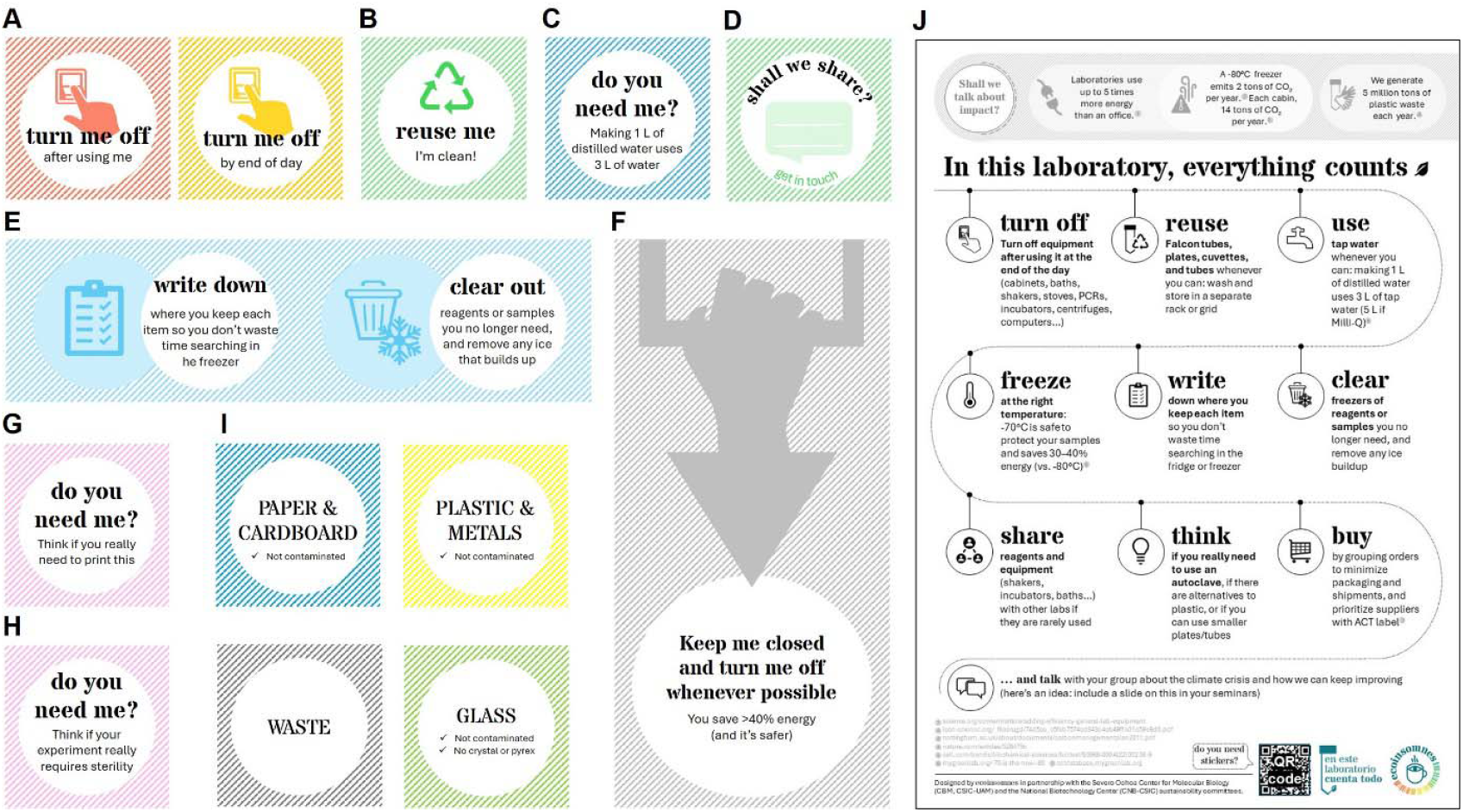
Stickers (A–I) and infographic (J) designed for the pilot scheme (English translation) (A) Stickers for electrical devices/equipment. (B) Sticker for trays/boxes with washed plastic cuvettes, tubes, and dishes that may be reused. (C) Sticker for distilled or milli-Q water taps/tanks. (D) Sticker for reagents or small equipment that a participant may choose to share with others. (E) Stickers for freezers. (F) Sticker for fume hoods. (G–H) Stickers for printers and autoclaves. (I) Stickers for recycling bins. (J) A4-sized summary infographic with key measures.

Second, we created a summary infographic sheet that reminded staff of key facts around the environmental impact of research and highlighted good practices; this included most items covered in the stickers and some additional points around procurement and encouraging discussions with colleagues around this issue (**Fig. 1J**). Print-ready Spanish versions of the stickers and the infographic are available from the corresponding author on request.

Third, the scheme was launched in both centres with a ~40-min live seminar that summarised key facts around the climate crisis and the environmental impact of research, reviewed previous schemes abroad, highlighted good practices and practical tips, and encouraged attendees to participate in the pilot scheme. Seminar slides are available from the corresponding author on request. At the end of the seminar, a ~20-min open session allowed attendees to discuss the seminar contents as a group, reflect on their colleagues’ interest and behaviours toward sustainability, as well as to provide suggestions for further actions.

Lastly, on the day of the seminar, a 30-second talking-head video summarising the scheme and promoting participation was distributed through CBM/CNB social media profiles.

### Data collection and analysis

The scheme was launched in January 2025. After the seminar, staff in both centres who decided to participate were invited to complete an online survey (~5 min) to capture their current behaviours. Each participating laboratory, service, or department (hereinafter referred to as “participant”) was represented by one staff member; as such, survey respondents were asked to consider behaviours in their laboratory/group rather than only their own when completing the survey. The relevant sustainability committee (CBM or CNB) subsequently provided respondents in their centre with a pack with stickers and infographic for their laboratory/group. To assess any changes produced in their laboratories/groups since the start of the pilot study, respondents were contacted via email and asked to complete a follow-up online survey (~10 min) in June 2025. Respondents were not sent a copy of their initial responses or reminded of them at the time of completing the follow-up survey. A copy of both surveys is provided in the **Electronic Supplementary Material**. At the start of each survey, respondents consented for their anonymised responses to be used for analysis and dissemination purposes. No economic compensation was provided.

Only data from participants who completed both surveys were used for analyses. The authors’ own groups also took part in the pilot scheme, but they were excluded from analyses to avoid bias.

Quantitative results were summarised using descriptive statistics (percentages, mean, standard deviation [SD], median, and range). Percentages were calculated using the relevant sample size for each question as a denominator, as not all questions were applicable to, or answered by, all participants. Paired (baseline vs. follow-up) responses were compared using the Wilcoxon matched-pairs test, a non-parametric test; two-tailed *p*-values are reported. Qualitative insight and participant quotes provided reflect comments entered as free-form text.

## Results

### Participants

189 groups (CBM: n=97; CNB: n=92) were targeted, of which 50 (CBM: n=32; CNB: n=18) accepted the invitation to participate (**Fig. 2A**). Some were lost to follow-up, so the final sample for analysis included 34 groups (CBM: n=21; CNB: n=13) (**Fig. 2B**). In the final sample for analysis, most participating groups were represented by postdoctoral or PhD researchers (n=9; 26% each), followed by technicians (n=8; 24%) and principal investigators (PI; n=3; 9%). Five participants (n=5;15%) were represented by staff in other roles (**Fig. 2C**).

**Fig. 2.**
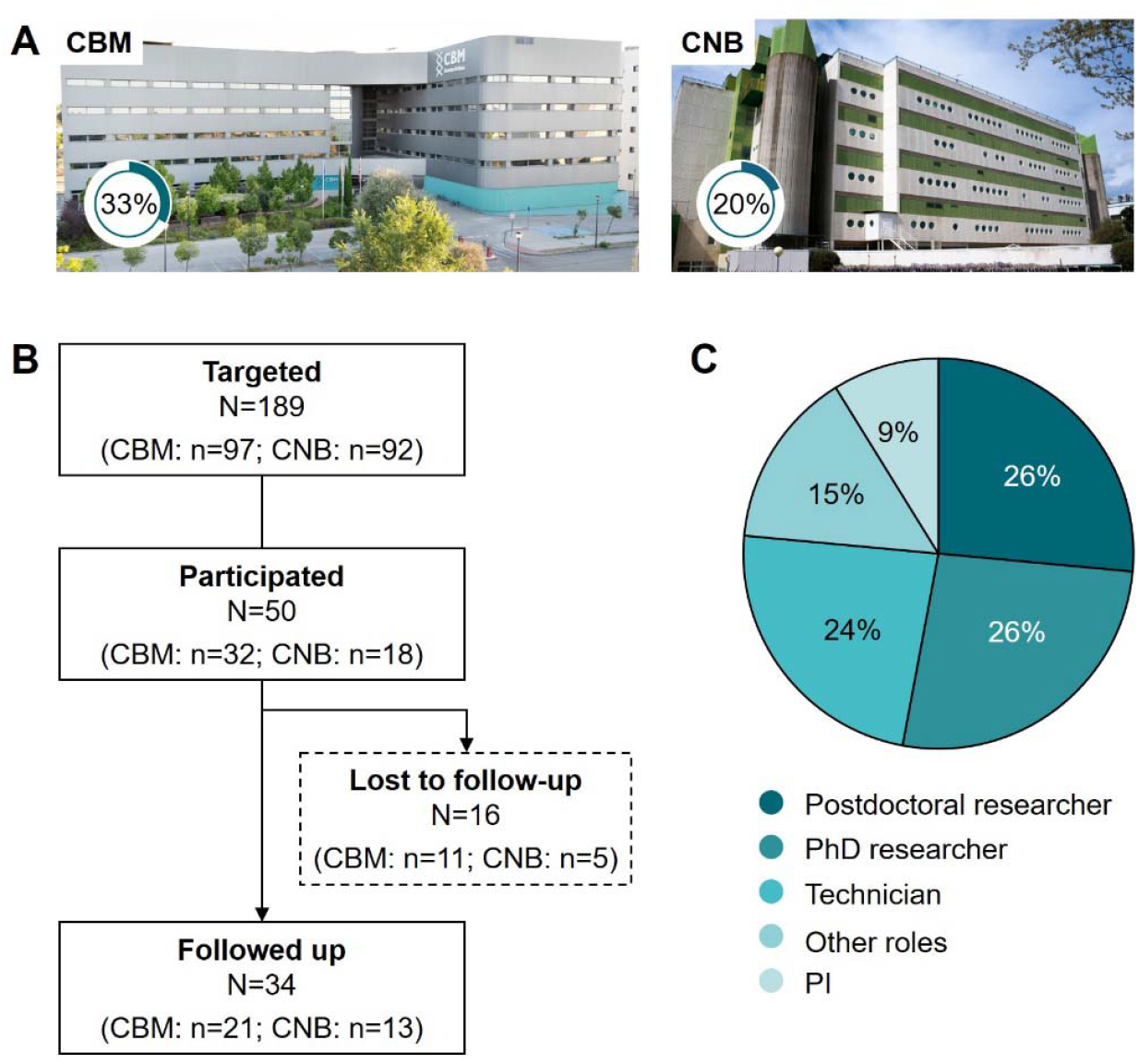
Participant disposition. (A) Images of both centres with percentages of targeted laboratories/groups who participated in the pilot study. (B Overview of participant enrolment; analyses used only data from the N=34 who were followed up for five months. (C) Distribution of staff representing each participating laboratory/group in this study; one person served as proxy for each participant. CBM, Severo Ochoa Molecular Biology Centre; CNB, Spanish National Centre for Biotechnology; PI, principal investigator.

### Energy use

#### Equipment switch-off

Participants were encouraged to switch off equipment after use or by the end of each day, when possible. At baseline, most participants already had this habit (mean of 4.2 [SD: 1.0]; scale: 1=“no equipment” to 5=“all equipment”). After 5 months, 10/34 participants (29%) had increased their rating, and the average score increased to a mean of 4.4 (SD: 0.7), although this was not statistically significant (*p*=0.23) (**Fig. 3A**). Two participants mentioned challenges with the equipment in communal areas. One (CNB, Other role) said “in our own lab, the equipment is turned off, but in shared areas like the culture room, responsibility becomes diluted and the same level of attention is not given.”

**Fig. 3.**
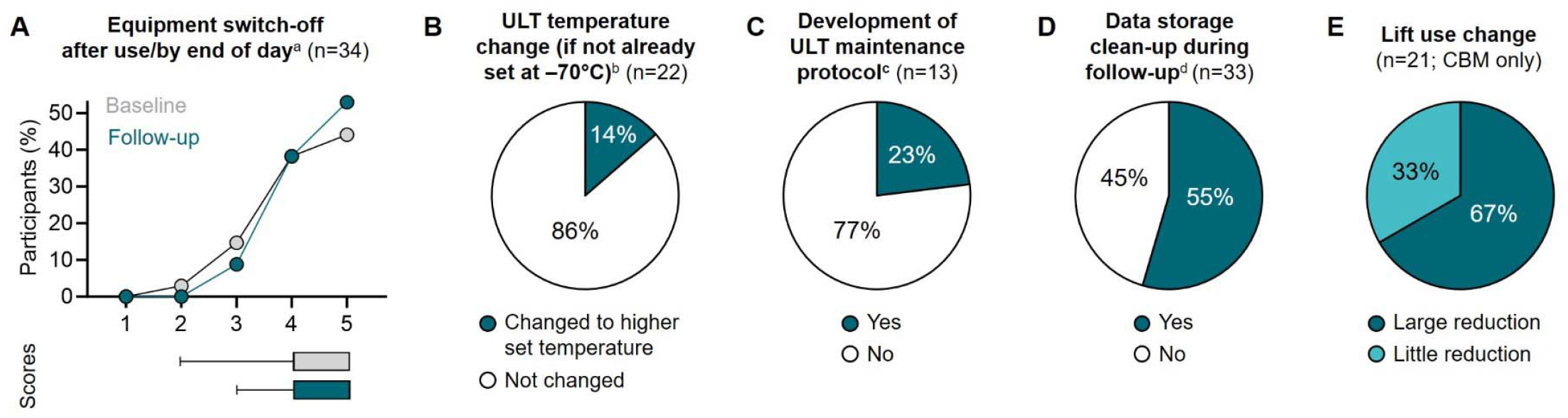
Changes in energy use. (A) Equipment switch-off (scale: 1=“no equipment” to 5=“all equipment”). Horizontal bars indicate score distribution (box and whiskers); *p*=0.23. (B) Temperature change of ULT freezers not already set at –70°C at baseline. (C) ULT maintenance. (D) Data clean-up. (E) Lift use. ^a^Survey question: “Do you turn off the equipment after using it or at the end of the day (shakers, incubators, water baths, centrifuges, PCR machines, ovens, computers…)?” ^b^Only responses from participants who had/used an ULT freezer, reported its set temperature at baseline and whether they had actively changed it during follow-up, and did not already have the ULT freezer set at –70°C at baseline were included for this analysis. ^c^Survey question: “Do you have any protocols for organizing or cleaning the ULT freezers?” Only responses from participants who responded no at baseline were included for this analysis. ^d^Survey question: “Since January, have you deleted old files from the servers to reduce energy consumption?” CBM, Severo Ochoa Molecular Biology Centre; PCR, polymerase chain reaction; ULT, ultra-low temperature.

#### ULT freezer temperature-setting and operation

Participants were informed of the energy saving that can be achieved setting ULT freezers to higher (less cold) temperatures without compromising sample integrity. At baseline, the participants’ ULT freezers were at a mean of –73.9°C (SD: 3.5°C); 8/30 (27%) of them were already set at –70°C (the target temperature this scheme recommended). Within the 5-month follow-up period, three of the remaining 22 (14%) had changed to a higher set temperature (**Fig. 3B**). However, none of the five ULT freezers that were set at –80°C at baseline had changed their temperature.

When asked about ULT freezer maintenance, decluttering, and sample organization, 43% already had measures in place at baseline. Within the 5-month follow-up period, 3/13 (23%) of those that did not have a protocol at baseline had put measures in place (**Fig. 3C**). Specific measures mentioned by participants included defrosting and removing ice build-up (n=7), general cleaning (n=5), and keeping an updated inventory/freezer map (n=2). Two participants mentioned that maintenance of shared freezers was a challenge.

#### Data storage

The seminar reminded participants to regularly remove stored data files that they may no longer require to save energy, even though there were no specific stickers for this. Within the 5-month follow-up period, 18/33 (55%) reported deleting old files from servers to save energy (**Fig. 3D**).

#### Lift use

Finally, in CBM (a seven-story building), stickers were placed on lifts and service lifts to encourage use of stairs when possible. All the participants in this centre reported at least some reduction in their lift use during the follow-up period. Most participants had reduced their use “a lot” (14/21; 67%) and some had reduced it “little” (7/21; 33%) (**Fig. 3E**). CNB participants were not asked about this because CNB lifts already had stickers in place before this pilot scheme.

### Water use

As a key measure to save water, participants were encouraged to use, when possible, tap water or distilled water rather than milli-Q water. At baseline, many participants already had this habit (mean of 3.3 [SD: 1.3]; scale: 1=“never” to 5=“always”). After 5 months, 16/33 participants (48%) had increased their rating, and the average score increased to a mean of 4.0 (SD: 1.1), which represented a statistically significant change (*p*=0.001) (**Fig. 4**). One participant (CNB, Postdoctoral researcher) described their practice in the follow-up survey as “first wash with tap water, followed always by distilled water to rinse.” However, three participants mentioned finding it difficult to convince others to adjust their protocols.

**Fig. 4.**
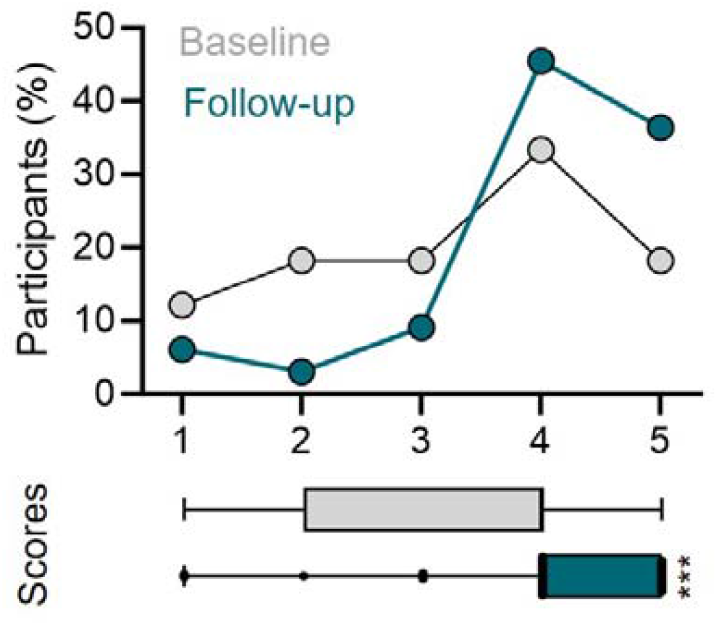
Changes in the use of distilled water. Survey question: “Do you use tap water (instead of distilled or milli-Q) whenever possible?” (scale: 1=“never” to 5=“always”). Horizontal bars indicate score distribution (box and whiskers); \*\*\**p* = 0.001.

Of note, three CNB participants reported issues with their centre’s water system resulting in cloudy and unreliable water. A subgroup analysis motivated by this revealed that mean ratings were higher for CBM than CNB participants, both at baseline (3.8 [SD: 1.1] vs. 2.4 [1.2]) and after follow-up (4.2 [SD: 0.7] vs. 3.6 [SD: 1.5]). However, CNB participants showed a clear improvement during the study, with a 1.3-point increase in their mean rating compared with a 0.5-point increase for CBM participants.

### Plastic use and sharing of reagents or equipment

Participants were encouraged to wash and reuse commonly used plastic containers when possible as a key measure to reduce plastic waste. At baseline, many participants already had this habit (mean of 3.2 [SD: 1.3]; scale: 1=“never” to 5=“always”). After 5 months, 16/33 participants (48%) had increased their rating, and the average score increased to a mean of 3.5 (SD: 1.3), although this change was not statistically significant (*p*=0.25) (**Fig. 5**).

**Fig. 5.**
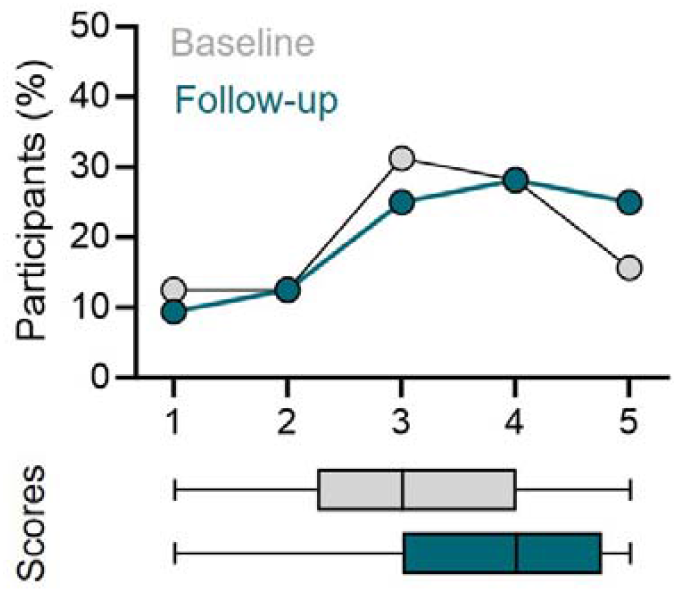
Changes in the reuse of plastic items. Survey question: “Do you reuse Falcon tubes, plates, cuvettes, or tubes whenever possible?” (scale: 1=“never” to 5=“always”). Horizontal bars indicate score distribution (box and whiskers); *p*=0.25.

Participants were also encouraged to share reagents or equipment with other laboratories, which can represent another effective way to reduce waste of plastic and other materials, as well as to save energy. Sharing with other laboratories was already a common behaviour prior to this scheme, with 25/31 (81%) practicing it at baseline. During the follow-up period, only two participants used stickers to label reagents or equipment for sharing with other laboratories. Another two participants noted that they had not used stickers but have offered sharing to other groups more informally. Six participants explicitly mentioned sharing reagents, including culture media (n=2) and isopentane (n=1). Of note, many more (n=18) explicitly mentioned sharing small equipment, including shakers (n=13), centrifuges (n=9), incubators/ovens (n=8), and PCR machines (n=6). Other examples included dissecting microscopes, cryostats, magnets, and specialised equipment for cell lysis, electroporation, or electrophoresis (n=1 each). Additional measures suggested by participants included reusing “empty [old] bottles for waste storage” (e.g., of acids; CBM, PhD researcher) and having “a central storage for commonly used reagents” (CBM, Technician).

### Procurement

Measures around procurement focused on bundling external orders and checking the environmental impact factor of potential providers; both were covered by the seminar and the infographic.

Participants were encouraged to bundle external orders when possible to reduce packaging and shipping emissions. At baseline, many participants already had this habit (mean of 3.5 [SD: 1.3]; scale: 1=“never” to 5=“always”). After 5 months, 12/33 participants (36%) had increased their rating, and the average score increased to a mean of 3.8 (SD: 1.1), although this change was not statistically significant (*p*=0.13) (**Fig. 6A**). One participant (CBM, PI) suggested a system to bundle orders at a centre level: “For instance, placing a [centralised] order for the main providers once a week.”

**Fig. 6.**
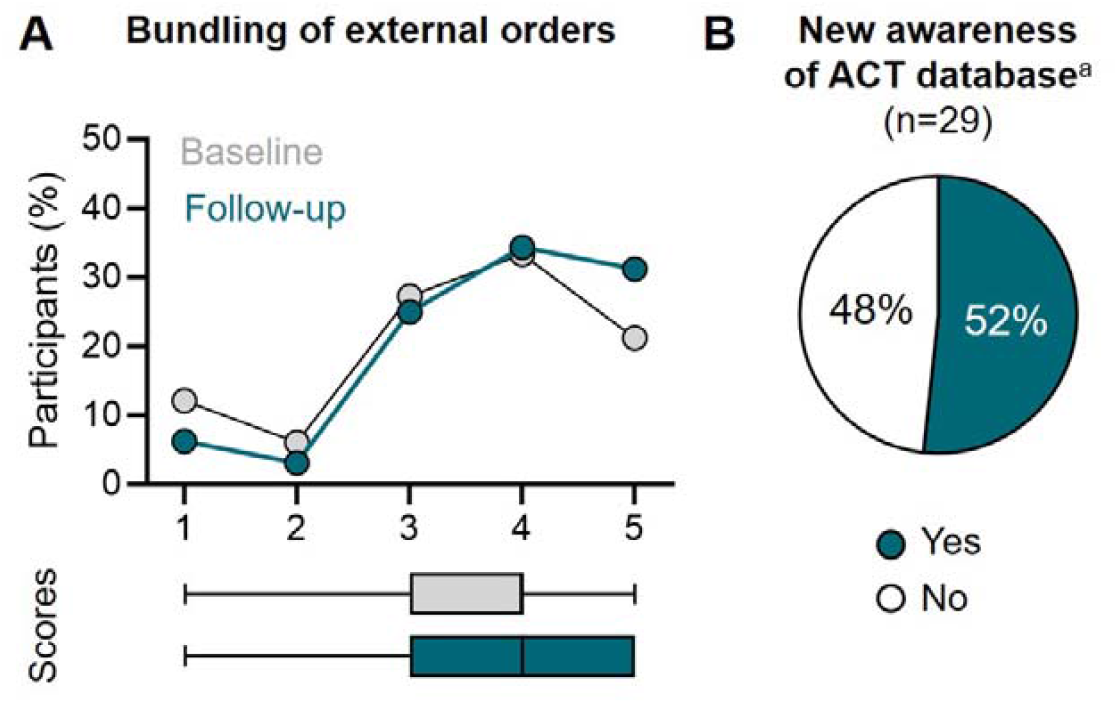
Changes in purchasing decisions. (A) Order bundling. Survey question: “Do you try to bundle external orders” (scale: 1=“never” to 5=“always”). Horizontal bars indicate score distribution (box and whiskers); *p*=0.13. (B) New awareness of ACT supplier database. ^a^Only responses from participants who were not aware at baseline were included for this analysis.

Participants were also informed about My Green Lab’s ACT database (19) and were encouraged to use it when choosing between different providers for external orders. Few participants (3/32; 9%) were aware of this database at the start of the pilot study. After the 5-month follow-up, 15 of the 29 participants that did not know the database at baseline (52%) were aware of it (**Fig. 6B**). During follow-up, three participants (including two who were not previously aware of this database) had used this database when preparing some orders.

### Colleague engagement

The seminar and the infographic highlighted the importance of discussing environmental sustainability as a group and encouraged researchers to engage their colleagues to make sustainability part of their group’s and centre’s culture.

At baseline, most participants (29/33 responses; 88%) had previously discussed sustainability, mainly in short or informal conversations, while 4/33 (12%) had not. Remarkably, as many as 31/33 (94%) discussed sustainability with their groups during the 5-month follow-up period alone, including 25 who discussed it in short/informal conversations and nine who discussed it in a seminar or group meeting.

Only two of the participants (6%) did not discuss sustainability with their colleagues during the follow-up period (**Table 1**).

**Table 1.** Colleague discussions on research sustainability.

| Have you discussed sustainability in the context of research in your laboratory/service? | Ever before baseline<br>(n=33) | During follow-up period alone<br>(n=33) |
| --- | --- | --- |
| Yes | 29 (88%) | 31 (94%) |
| In short/informal conversations<sup>a</sup> | 24 (73%) | 25 (76%) |
| As a group (in meetings or seminars)<sup>a</sup> | 8 (24%) | 9 (27%) |
| No | 4 (12%) | 2 (6%) |

<sup>a</sup>Answers were not mutually exclusive.

Overall, some participants (n=6) mentioned the measures were well received by their colleagues, while others (n=5) reported poor interest/response. One participant (CBM, PI) summarised the situation saying that “we are all aware, but sustainability measures are not always put into practice.” Another (CBM, PI) noted that changing habits is difficult: “In the end, it’s usually the people who were already doing it before who continue to do it. For those who weren’t, it’s still hard for them to apply these changes unless they’re constantly reminded.”

### Overall assessment

Participants generally found this pilot scheme useful. When asked about its usefulness, the median rating was 3 or “fairly useful” (scale: 1=“not useful at all” to 5=“indispensable”; n=34). When asked to rate the individual elements separately, the seminar was the most valued (median rating: 4 or “very useful”; n=21 [13 participants did not attend]) followed by the infographic and stickers (both with a median rating: 3 or “fairly useful”; n=33 for the infographic [one participant did not use it] and n=34 for stickers) (**Fig. 7**). Two participants spontaneously identified the high level of visual noise in laboratories as a barrier to the effectiveness of stickers.

**Fig. 7.**
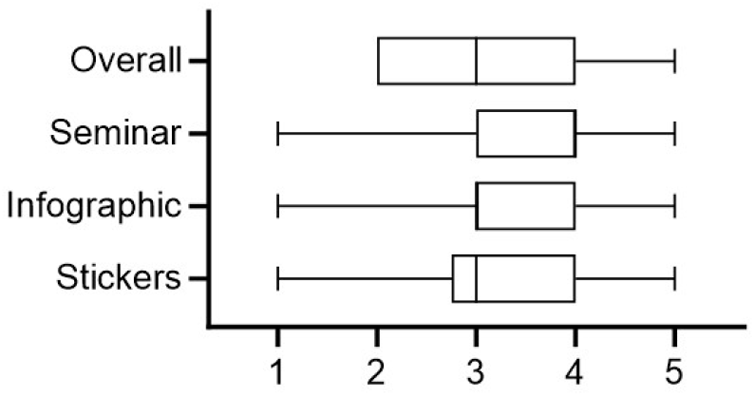
Usefulness of scheme (overall and by component) Survey question: “In your opinion, how useful has the scheme been overall [/has each component of the scheme been]?” (scale: 1=“not useful at all” to 5=“indispensable”). Horizontal bars indicate score distribution (box and whiskers).

In terms of future steps, five participants suggested regular reminders/seminars and even brief presentations about sustainability in individual laboratories, two suggested emphasising the economic savings that can be achieved with energy measures or through sharing reagents, and one suggested competitions/challenges with rewards. Of note, one participant (CNB, PhD researcher) stated that “it would be good to encourage PIs from various laboratories to attend the initial seminar… many people who attended the seminar were unable to put stickers in their respective laboratories because they were not allowed to do so.” Participants also highlighted the need for wider leadership to take action, including one (CBM, Technician) who suggested “institutional regulations” and called on CSIC to get involved in sustainability efforts, and another (CNB, PI) who requested more efficient heating control for the building.

## Discussion

Raising awareness of practical strategies to reduce the carbon footprint of laboratory research is becoming increasingly important. This pilot scheme engaged 50 laboratories, scientific services, and departments across two of the largest biomedical research centres in Spain. It developed a scalable, visual-oriented communications strategy designed with input from research centre staff, which included a live seminar and a large but coherent set of graphic materials. Pilot results provide a breadth of quantitative and qualitative insights that can guide further efforts and escalation at a regional and national level.

Our results indicate engagement in sustainability-related behaviours within five months; for instance, nearly a quarter of participants developed measures for energy-efficient operation of freezers, over half deleted old data from storage to save energy, and everyone reduced their use of lifts. Behaviours regarding equipment switch-off, reuse of plastic items, prioritization of tap water when possible, and bundling of external orders all shifted towards best practice.

The findings also reinforce the widely accepted view that sharing is a core principle of scientific culture; one that promotes efficiency, reduces waste, and strengthens collaboration between laboratories. Of note, this scheme has motivated the creation of an online reagent database at CBM, which will allow researchers to list reagents they are willing to share with others. However, the management/maintenance of shared equipment was a key challenge raised by researchers and the adoption of some sustainable practices was met with resistance. For instance, while three participants raised the temperature of their ULT freezers, most did not. In particular, all ULT freezers set at –80°C (which requires much more energy) remained at that temperature. Similarly, some participants mentioned challenges convincing colleagues that modified protocols (e.g., using different water sources or reused reagents or containers) would still produce reliable results. These findings highlight the value of sharing experiences with modified protocols to build trust in their reliability among colleagues. Pertinent to that, this scheme prompted almost all participants to engage in both informal and formal conversations about sustainable research practices. Participants also rated the initial seminar session very positively, further indicating the key role of face-to-face communication. Combining introductory sessions with regular follow-ups may enhance impact and long-term engagement, and the open exchange of ideas and concerns can contribute to resolving issues. For instance, although participants in this study significantly improved their use of water resources in this study, the data revealed concerns among CNB participants regarding the quality/reliability of tap water. This underscores the importance of effective communication channels between sustainability committees and the management and leadership teams in each centre, who should act to address concerns as appropriate. While volunteer-led sustainability committees have been instrumental in initiating changes in different areas, their work is often constrained by factors including limited time, continuity, and specialist training. The appointment of dedicated sustainability officers can fill these gaps by overseeing and optimising the use of shared resources; serving as an expert point of contact to channel questions, suggestions, or concerns; and connecting researchers with their local sustainability committees and other relevant resources. Therefore, introducing this role could dramatically enhance the long-term effectiveness of sustainability efforts (23–25).

Overall, these results highlight the need to make environmental sustainability part of the wider laboratory and organisational culture. Participation (the ratio of participants to target sample) was higher at CBM compared to CNB, which may be partly due to each group having a sustainability point of contact at CBM, but not at CNB. In terms of roles, few PIs were directly engaged in this pilot study (only 9% of participants). Engaging senior staff (e.g., senior postdocs and PIs) and institutional leadership teams, in addition to funders and other stakeholders, can significantly enhance engagement (26) by supporting the involvement of more junior researchers and by sending a strong message of institutional commitment (23). In this regard, CBM’s sustainability committee is developing a programme that integrates specific training on sustainable laboratory practices into the onboarding process for all new members, with a focus on early-career researchers to encourage environmentally responsible behaviours from the start of their career. Where appropriate, leadership teams must offer recognition and incentives for sustainable practices, while also addressing counterproductive behaviours, including those that may arise from PIs.

Embedding sustainability into codes of conduct and organisations’ core values may be helpful first steps towards this end. Research centres may also set up competitions and awards to recognise role models and showcase best practices, and CSIC could introduce a nationwide sustainability accreditation or badge, which would work similarly to its equality badge.

We acknowledge that this pilot study presents some limitations. For instance, participants joined the study voluntarily and were therefore likely to place more value on sustainability than respondents chosen at random would (potential sampling bias). While behaviours within the laboratory are arguably best assessed by researchers themselves, many analyses were descriptive; since it was not possible to obtain data on energy or water use from the centres’ management, changes in consumption could not be measured in this pilot study. Further, although our 5-month follow-up successfully identified shifts towards positive behaviours, it is uncertain whether behaviours will gradually improve or revert over time. Finally, actions for leadership were intentionally out of scope for this pilot, although these are absolutely required to promote and complement bottom-up efforts towards sustainability.

Pilot schemes elsewhere have been successful in securing subsequent large-scale rollout of measures where there was leadership/organisational support (23). Consistent with previous research (10), our results show a considerable baseline interest in sustainability among researchers, suggesting there would be strong support for putting CSIC’s recently unveiled sustainability action plan into practice across the Spanish research community. However, this will require concrete commitments from CSIC towards the funding and timely implementation of a coherent set of actions. In this context, the present pilot demonstrates practical, scalable strategies and actions that can be readily scaled up and integrated into publicly-led, nation-wide schemes.

## Conclusion

We developed a scalable, visual-oriented communications strategy that was successfully piloted across two research centres in Spain. The scheme promoted sustainable behaviours within research laboratories, achieving engagement and shifts towards best practice across metrics. However, the findings also revealed key areas for improvement. These insights could guide broader institutional strategies aimed at enhancing sustainability in research.

## Data availability

All data supporting this article have been included as part of the submission. Response data from individual participants cannot be made available due to consent restrictions. Print-ready Spanish versions of the developed materials are available from the corresponding author (PI) on request.

## Author contributions

All authors contributed to study concept and design; RMF, SA, CC, MLS, and PI contributed to data collection; PI analysed the data and drafted the initial manuscript draft; and all authors contributed to data interpretation, edited the manuscript, and approved the final version for submission.

## Acknowledgements

The authors thank all CBM and CNB staff who support sustainability efforts, the members of their respective sustainability committees, and especially all participants of this pilot study for their involvement. Bruno Martín de la Llama and Mario Juárez (ecoinsomnes) provided feedback on the overall communications plan and on initial versions of the materials developed. María José Martín Pereira (CBM) and Jennifer Palencia Ortega and Susana de Lucas (CNB) supported internal communications at pilot launch and provided the building photographs included in Fig. 2.

## Electronic Supplementary Material

### Survey 1 (baseline)

| Question | Answer options |
| --- | --- |
| 1. Do you turn off the equipment after using it or at the end of the day (shakers, incubators, water baths, centrifuges, PCR machines, ovens, computers...)? | 1 (No equipment) / 2 / 3 / 4 / 5 (All) |
| 2. Do you reuse Falcon tubes, plates, cuvettes, or tubes whenever possible? | 1 (Never) / 2 / 3 / 4 / 5 (Always) |
| 3. Do you use tap water (instead of distilled or milli-Q) whenever possible? | 1 (Never) / 2 / 3 / 4 / 5 (Always) |
| 4. What temperature (°C) is your ultra-low temperature freezer set to? | [Free-text response] |
| 5. Do you have any protocols for organising or cleaning the ultra-low temperature freezers? | [Free-text response] |
| 6. Do you share any reagents or small, less-used equipment (rotators, shakers, etc.) with other laboratories/services? Please indicate which ones. | [Free-text response] |
| 7. Do you try to bundle external orders? | 1 (Never) / 2 / 3 / 4 / 5 (Always) |
| 8. Were you **already** familiar with the ACT label [hyperlink: https://act.mygreenlab.org/] for suppliers? | Yes / No |
| 9. Have you discussed sustainability in the context of research within your laboratory/service? | [Respondents could select one or more answers from:]<br>• Yes: as a group (in meetings or seminars)<br>• Yes: in informal/short conversations<br>• No |
| 10. Have you discussed sustainability in the context of research within your laboratory/service? Can you think of any measures you could implement in your laboratory/service to become more sustainable? | [Free-text response] |

### Survey 2 (follow-up)

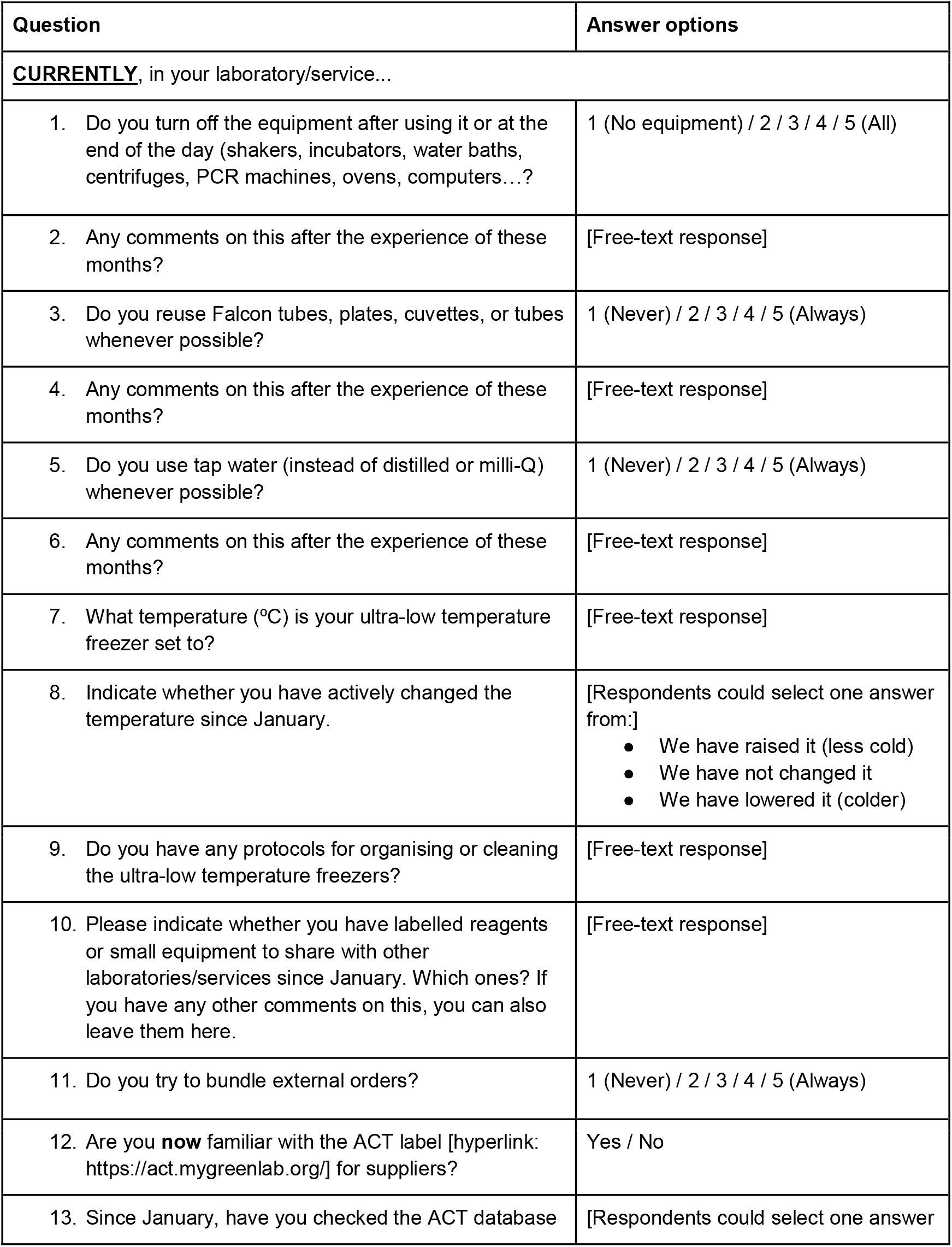
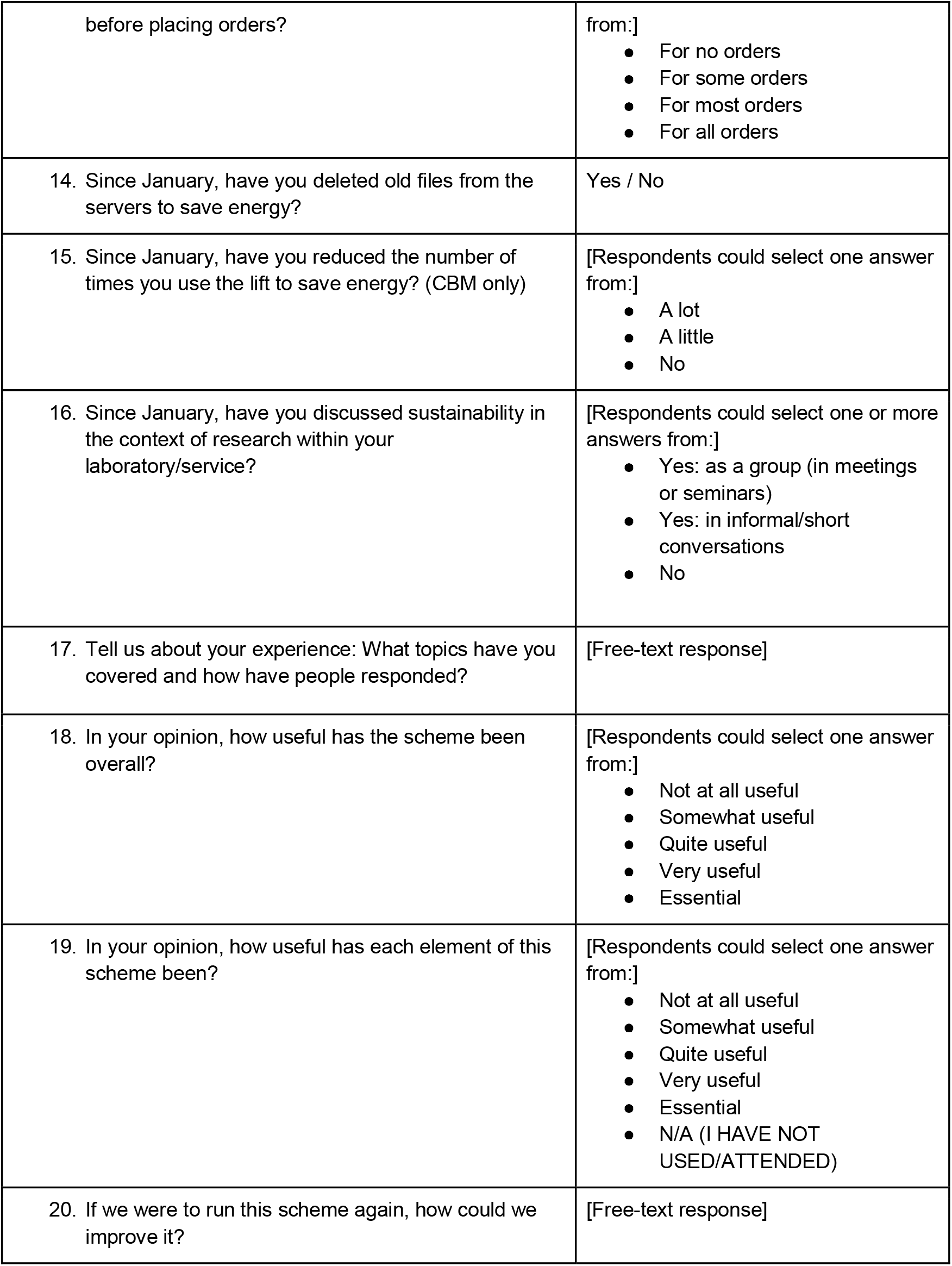

